# How Turkey vultures tune their airspeed to environmental and behavioral factors

**DOI:** 10.1101/2022.07.08.499337

**Authors:** Jonathan Rader, Tyson L. Hedrick

## Abstract

Animals must tune their physical performance to changing environmental conditions, and the breadth of environmental tolerance may contribute to delineating the species’ geographic range. A common environmental challenge that flying animals face is the reduction of air density at high elevation and a reduction in the effectiveness of lift production that accompanies it. Turkey vultures (*Cathartes aura*) inhabit a >3000 m elevation range, and fly considerably higher, necessitating that they compensate for air density differences through behavior, physiology, or biomechanics. We predicted that birds flying at high elevation would demonstrate higher median flight speeds while maintaining similar glide angles. We used 3-dimensional videography to track Turkey vultures flying at three elevations and found a negative relationship between median airspeed and air density that matched our prediction. Additionally, neither the ratio of horizontal speed to sinking speed nor flapping behavior varied with air density. These results were robust to varying flight behavior (climbing vs. level flight). Finally, we derived a glide polar from the free-flying vultures and showed that they are proficient at tuning their flight speed to minimize their cost of transport during straight-line flight, but transition to a minimum power strategy during gliding turns.

## Introduction

The breadth of environmental conditions that species tolerate and exploit may, in large part, determine their geographic extent. Intuitively, species that can tolerate only a narrow set of environmental variables may be expected to have smaller geographic distributions than species that thrive in more diverse conditions [1]. Thus, exploring how species whose ranges span broad environmental gradients or inhabit variable environments compensate for environmental challenges, and what prevents others from reacting similarly, may shed light on how geographic range is constrained.

Life at high elevation, and the correspondingly reduced air density, presents a two-fold challenge to locomotor performance in flying animals [2]. First, a physiological challenge of low air density stems from the reduction of available oxygen for respiration [2,3], which can lead to hypoxia and decreased metabolic power output. Secondly, reduced air density poses a physical challenge to fliers, as it decreases the effectiveness of lift generation [2,4]. However, birds can be found at the highest elevations [5,6], suggesting that some species are able to compensate for these hardships. A variety of cardiopulmonary adaptations allow high flying birds, such as bar-headed geese [5,7] and several lineages of Andean waterfowl [6,8], to obtain the oxygen that they need for aerobic respiration. There are a variety of mechanisms that fliers can use to compensate for the aerodynamic consequences of flight in low air density. As was documented in tropical hummingbirds by Feinsinger et al. [9], high elevation species tend to have larger wings relative to their body mass than their low-elevation counterparts. Additionally, birds can adapt to flight in low air density by increasing power output [9], flapping more than they would in higher density air [10], or by using higher amplitude wing strokes [11]. One question that has received less attention, though, is whether birds are capable of compensating for reduced air density at high elevation behaviorally, such as through modulation of airspeed. While not applicable to hovering flight or to landing and takeoff, modulation of airspeed may otherwise offer a low to no cost method for maintaining flight performance across air density gradients. Modern techniques that facilitate high-resolution tracking of birds in the field, either via GPS technology [e.g.: 12], or by high-definition videography [13,14], can be used to look for differences in flight speed among bird populations living at different elevations.

Bird species with broad geographic ranges may be exposed to large elevation gradients, and this provides an opportunity to study how they tune their locomotor performance to different environmental conditions. Turkey vultures (*Cathartes aura*) are common throughout North America, inhabit an elevation range of >3000 m [15], and have been reported flying at much higher altitude [16–18]. No evidence is available to suggest that any morphological differences in wing area or flight muscle mass exist among *C. aura* throughout their range, though this has not been addressed explicitly. However, they could alter their power output via increased flapping frequency or higher amplitude wing strokes when challenged by low air densities. Vultures are primarily gliding fliers [19–22], and could also adopt a strategy where they maintain the same true airspeed (which is the same as ground speed in still air) by increasing their glide angle, effectively increasing their power output by cashing in potential energy at a greater rate. Furthermore, *C. aura* consume almost exclusively carrion, which is a food resource that is sparsely distributed and highly ephemeral [23,24]. Thus, it would seem advantageous for *C. aura* to minimize their energetic expenditure while foraging, and suggests that increased flight power output is an unlikely adaptation to high elevation life. Thus, we hypothesized that vultures accommodate flight at varying air densities by changing their true (i.e. observed) airspeed, increasing it as air density decreases. Mathematically, this is identical to the vultures maintaining the same equivalent airspeed (i.e. airspeed corrected for air density) at all air density conditions they experience. We further predicted that vultures would avoid activities that increase their cost of flight: that they would not increase flapping in low air density, and that they would maintain similar glide angles throughout the elevation range.

We addressed these hypotheses by recording vultures flying at three sites along a ∼2000 m elevation gradient to examine how they tune their flight performance to compensate for the air density gradient. As expected, and regardless of the recorded vulture flight behavior, there was a negative relationship between observed airspeed and air density. Vultures did not flap more frequently in lower density air, and the observed change in speed was a near perfect match to that theoretically required to maintain lift, suggesting that other factors such as wing morphing are also unimportant in this case. This study demonstrates how field studies can illuminate the relationship between biomechanical performance and ecology.

## Methods

### Vulture recordings

We recorded vultures returning to roost sites on 22 separate afternoons in May, June, and July 2015, and September 2016 at three locations: the Orange County Landfill, Chapel Hill, NC, USA (35°58’9.23”N, 79° 4’54.71”W), the University of Wyoming campus, Laramie, WY, USA (41°18’44.15”N, 105°35’1.04”W, see Fig. 1) and the Alcova Lakeside Marina, Alcova, WY, USA (42°31’45.99”N, 106°46’44.21”W). Each roost colony was comprised by >50 individuals. We were unable to identify and track identities of individual bird and birds readily transited into and out of the camera field of view multiple times during recording bouts, so to pseudoreplication of these individuals, the resulting data were primarily analyzed by reducing each recording session to the median airspeed of all birds in that recording. Subsequent analyses of airspeed were conducted on those medians. There were multiple recordings per day, however we treated each of these separately because ambient temperature and humidity change throughout the day, so air density also varied among recordings.

**Figure 1.**
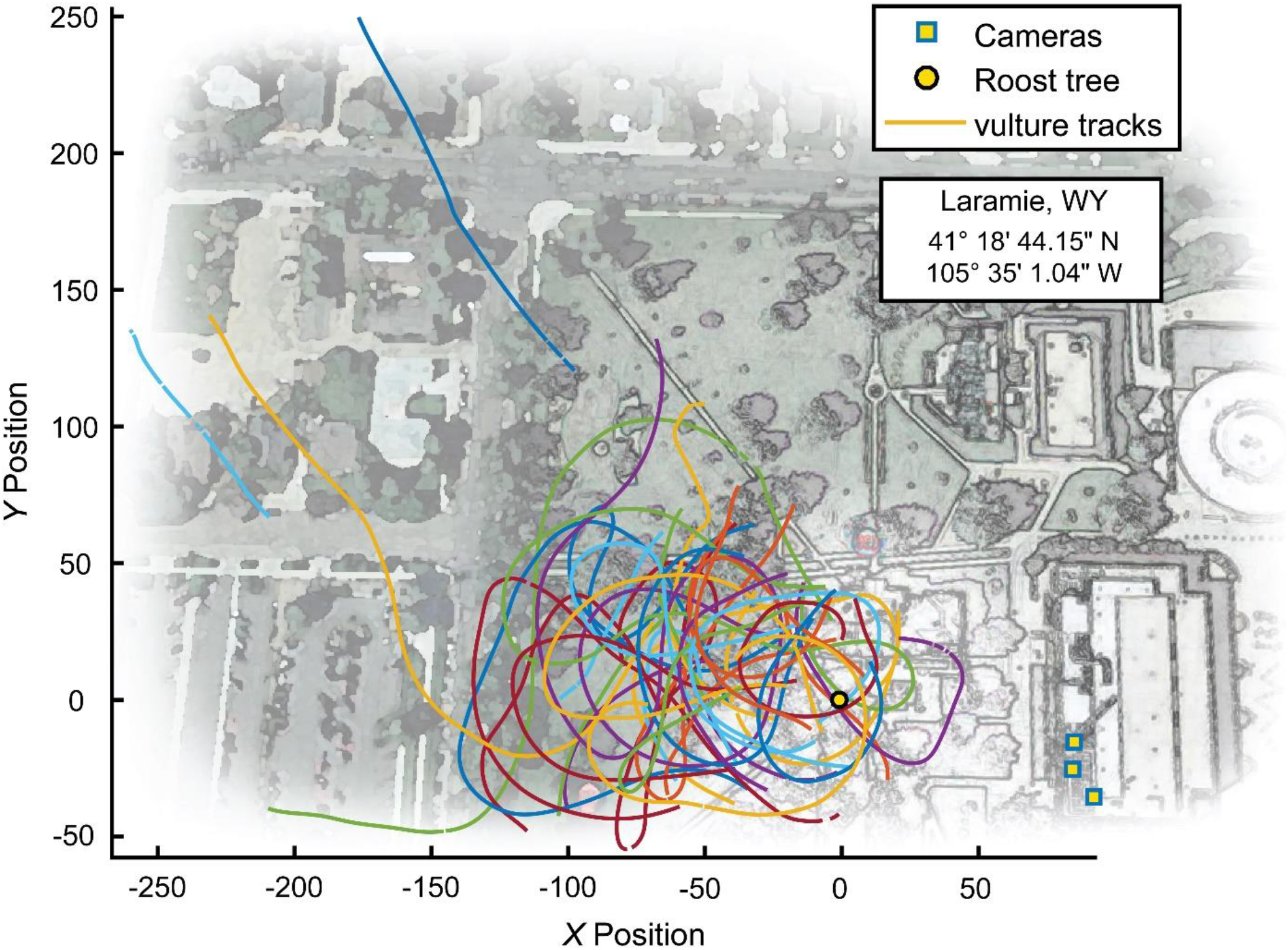
Overhead view of the Laramie, WY recording site. Blue squares denote camera locations atop the Biological Sciences building at the University of Wyoming, and the black circle shows the center of the roost trees. The multicolored tracks depict a sampling of the vulture tracks recorded from one recording bout.

Video data were collected with three digital SLR cameras (Canon OES 6d, Canon inc., Ōta, Tokyo, Japan) at 29.97 Hz and images had dimensions of 1920 by 1080 pixels. The cameras were arranged in a staggered setup, with intersecting views of the tops of roost trees and airspace above and around them (Fig. 1). Due to the altitude at which the vultures approached the roost trees, and the requisite upward angle of the cameras, we were unable to use a standard wand calibration [e.g.: 13,14]. Instead, we obtained a preliminary calibration using shared views of the flying vultures, digitized using the MATLAB (The MathWorks, Natick, MA, USA) package DLTdv5 [25]. After the initial calibration was complete, we automated tracking of the vultures using a computer vision workflow adapted from that used in Evangelista et al. [26]. In brief, birds were detected in each video file by using a 30-frame moving average background subtraction routine plus a fixed background mask for the trees. The resulting background-subtracted images were cleaned with an erosion-dilation operation. The [*u,v*] pixel coordinate of each remaining foreground object was recorded as a possible vulture detection. These 2D detections from the three cameras were combined to compute a 3D point by searching the possible 2D point combinations for ones that produced a 3D reconstruction residual of less than 3 pixels. Points generated from either two or three cameras were accepted. Once the sets of 3D points for all video frames were generated, we joined the resulting 2D+3D datasets across time using a set of Kalman filters to predict the expected position of the birds from frame *n* in frame *n*+1 and then a Hungarian assignment operation to match the observations in frame *n*+1 to these predicted positions. Unmatched observations started new tracks, and tracks with more than 20 missed detections in sequence were discontinued. Tracks with fewer than 300 data points were dropped from the dataset. The calibration was refined using the complete set of digitized points from the vulture tracks for each recording session, and we used the distances between the cameras to scale the scene. The cameras were aligned to gravity using their onboard roll-leveling feature, and we measured their pitch inclination using a digital inclinometer affixed to the hot-shoe mount on the top of the base camera body. Finally, the scene was oriented to a geographic frame of reference by aligning to the compass vector between the base camera and the roost tree.

Individual bird 3D tracks in the scaled and aligned dataset were smoothed using a zero-lag digital Butterworth low-pass filter with an 8 Hz cutoff frequency. Velocity vectors were calculated from this position time-series by fitting a quintic spline polynomial and differentiating it. We added the wind speed vector (see below) to this ground reference frame velocity vector to get each bird’s observed airspeed.

### Air density and airspeed

Wind and weather conditions during the recording sessions were recorded from the closest National Oceanic and Atmospheric Administration (NOAA) weather station for all locations, and from a rooftop-mounted weather station atop the University of Wyoming Biological Sciences building, adjacent to the roost. Because of the distance between the recording sites and the weather stations, and because ground-level wind conditions may not reflect the conditions experienced by the birds, we estimated the magnitude and direction of wind conditions during each recording session from the ground speeds of the birds as they flew in different directions, following the methods of Sherub et al. [27]. Absent any wind or other directional factors, bird ground speeds are not expected to vary with flight direction such that a plot of the two components of their horizontal velocity vector form a circle centered on [0,0]. A wind alters the center of the circle. For example, a 5 ms^−1^ wind in the +*X* direction moves the center to [5,0]. Thus, we estimated the wind speed and direction experienced by the vultures as the center point of a circle fit to the *X* and *Y* components of their measured groundspeeds over a recording session. This method depends on having a sample of flights headed in all compass directions. Consequently, we omitted trials with vulture tracks comprising less than 90% of the full circle. Ground speed was calculated as the first derivative of vulture position with respect to time (see above). Vulture airspeed was then obtained by subtracting the *X* and *Y* components of the estimated wind speed from their respective ground speed components of the vulture tracks. No sustained thermal soaring behavior was observed during the recording periods, and we were unable to assess vertical movement of the air, so we assumed that non-flapping tracks represented gliding flight.

Air density (*ρ*) was calculated from mean barometric pressure, ambient air temperature, and dew point readings from the NOAA weather stations during the recording periods using the formula for density of moist air from [28, pg. 37]:

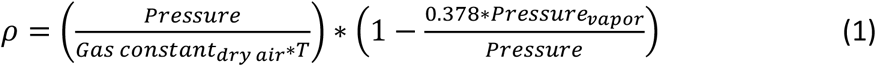

We calculated median airspeeds for each track, and from those the median airspeed of vultures during the entire recording session. Because we were unable to assign individual identifiers to birds and track them beyond the camera field of view, it is highly probable that individual vultures contributed multiple tracks to the dataset by appearing during successive recording sessions at the same site. However, these individual replicates would have been collected at separate points in time and thus represent flight under different conditions (e.g. air density, wind direction relative to flight direction, and even body mass).

We hypothesized that vulture airspeed would decrease as a function of air density (*ρ*) in the sample (predicted slope = -5.22) based on a linear approximation of the theoretical relationship between airspeed and *ρ* and the median airspeed of vultures at the maximum *ρ* in the sample (*ρ* = 1.227):

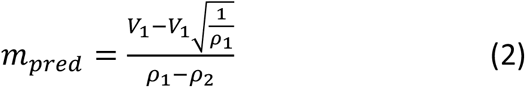

where *m*_*pred*_ is the predicted linear approximation slope, *ρ*_1_ is the highest air density in the sample, *ρ*_2_ is the lowest air density in the sample, and *V*_1_ is the median airspeed of vultures flying in the highest *ρ* conditions in the sample. The estimate of airspeed at *ρ*_2_ is based on equation 5 in Pennycuick [29]. Although the theoretical relationship between *V* and *ρ* is non-linear, because the range of *ρ* is small, a linear approximation provides a convenient and easily testable hypothesis.

We tested a set of multiple linear regression models to evaluate the relationship between median airspeed and *ρ* as well is the impact of wind speed on the relationship. We used AICc to select the best fit among these candidate models. Additionally, to assess whether flight behavior (i.e. climbing, descending, or level flight) might influence the relationship between air density and airspeed, we parsed the data into climbing, descending, and level flight tracks using vertical speed thresholds of (*V*_*Z*_,*med* > 0.25 m/s), (*V*_*Z,med*_ < -0.25 m/s), and (0.5 m/s > *V*_*Z,med*_ >-0.5 m/s) respectively, and used ordinary least-squares (OLS) regression to evaluate the relationship between *V* and *ρ* in each set. We used a Wald *χ*^2^ test to assess whether the model slopes differed from *m*_*pred*_.

### Air density and flapping behavior

The automatic tracking algorithm detects and tracks the visual centroid of the vultures in each video frame (as opposed to a fixed point on the body, such as the head). Because of this, and due to the large size of the birds’ wings, their tracks appear as a sinusoidal wave pattern when the birds flap, contrasting with comparatively smooth gliding tracks. We exploited this to assess whether the vultures flapped more in lower air density, a sign that they might be compensating for decreased lift by modulating power output. We used a custom MATLAB program to detect that characteristic sinusoidal track pattern and coded each video frame of each track with a binary (0) gliding, or (1) flapping. We then used the mean to quantify the proportion of the track the bird spent flapping vs. gliding. Further analyses of flapping behavior were restricted to tracks closer than 350 m from the cameras, as this appeared to be the maximum distance at which flapping is detectable (see Fig. 3). We also excluded tracks within 50 meters of the roost to avoid tracks in which the birds were making their final landing approach, which was characterized by a large amount of flapping not necessarily related to the density of the air. Data were again collapsed to medians for each recording bout. We used OLS regression to look for a relationship between incidence of flapping and *ρ*. Finally, we assessed whether the proportion of flapping in tracks using OLS regression on the mean, or the probability of detecting flapping (binary logistic regression) varied with wind speed.

### Glide polar calculation

Finally, we also constructed a glide polar from the vulture tracks by fitting the observations to the following equation [30]:

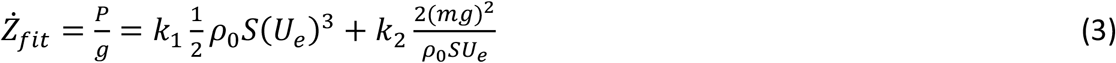

Where 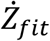 is the sinking rate that expends potential energy at the same rate as the overall kinematic power *P* (i.e. the instantaneous rate of change in kinetic and potential energy), *g* is the magnitude of gravitational acceleration, *k*_1_ is a coefficient for parasite and profile power, *ρ*_0_ is standard air density, *S* is wing area, *U*_*e*_ is equivalent airspeed (i.e. airspeed adjusted for air density), and *k*_2_ is the induced power coefficient. See Hedrick et al. (2018) for further details on this calculation. Literature values of 0.456 m^2^ and 2.18 kg [31] were used for *S* (wing area) and *m* (body mass), respectively. However, these values have no effect on the resulting glide polar, only on the numerical values for *k*_1_ and *k*_2_ arrived at by the statistical fitting process, a general linear model.

Reducing each vulture flight track to its median values, as was done for the air speed versus air density analysis results in a dataset with a compressed range of equivalent airspeeds whereas fitting a glide polar requires a wide range of airspeeds. Thus, for the glide polar calculations we created an alternative dataset by sampling individual data points from the entire set of flight tracks. In constructing this dataset, we first established the following criteria for possible inclusion: 1) non-flapping, 2) instantaneous turn radius > 20 m, 3) non-landing, 4) negative *P* (i.e. negative instantaneous summed rate of change in kinetic and potential energy), 5) P > 5^th^ percentile, and 6) distance to cameras < 350 m. From the resulting set of approximately 473,000 data points, we selected approximately 3000 equally spaced points, matching the number of vulture flight tracks.

For comparison with the dataset used to compute the glide polar, we produced a “gliding turns” dataset that might reflect brief thermal use or other aerial behaviour beyond straight gliding. For this dataset we used the following criteria: 1) non-flapping, 2) instantaneous turn radius < 15 m, 3) non-landing, 4) *P* in the 5^th^ to 95^th^ percentile range, and 5) distance to cameras < 350 m, the limit of flapping detection. From the resulting set of approximately 218,000 data points, we again selected approximately 3000 equally spaced points for inclusion in the dataset.

## Results

### Vulture tracks, air density, and flight speeds

We collected 3027 vulture flight tracks representing 18 hours of vulture flight time.Median vulture airspeeds, summarized by recording bout, ranged from 7.5 to 12.58 ms^−1^ with an overall median airspeed of 10.12 ± 0.87 ms^−1^. There was a large amount of variation in airspeed among vulture flight tracks in each recording session, median absolute deviations (MAD) ranged from 16% to 68% of the median (Fig. 2). After correcting for ambient temperature and relative humidity, *ρ* ranged from 0.890 to 1.227 kg m^−3^ (see Fig. 2).

**Figure 2.**
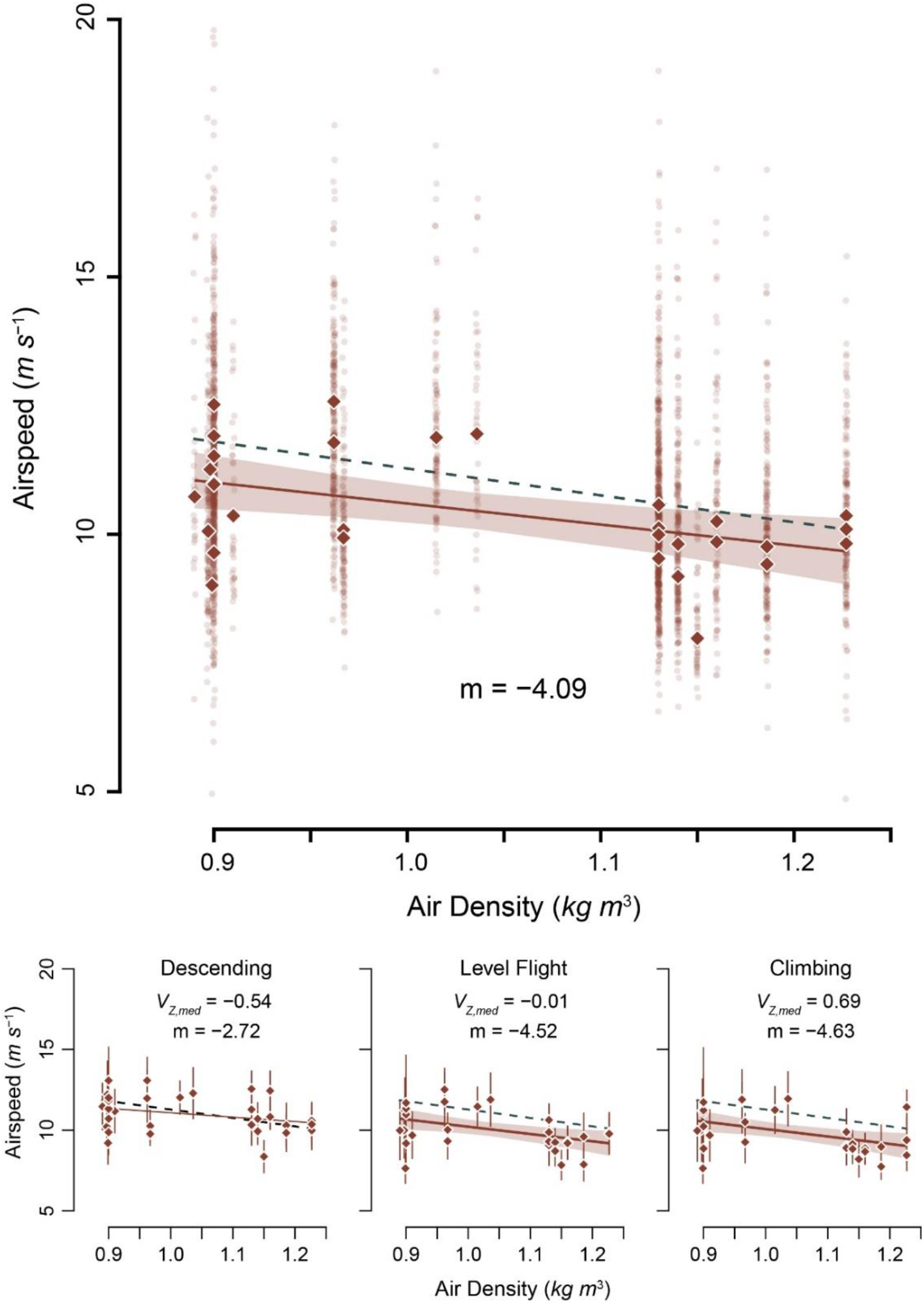
Median airspeed decreased with increasing air density. Transparent data points show median values for each track, highlighting the large variation in the sample. Analyses were conducted on median values for each recording session, depicted by solid diamonds. The dashed green line shows the predicted slope (−5.22), while the solid brown line shows the modeled slope with its 95% confidence interval (shaded region)

The best performing model, via AICc, included effects of *ρ* and wind speed (AICc weight = 0.67). Median vulture airspeed (*V*_*med*_) decreased with *ρ* (estimated slope *m*_*ρ*_ = -3.73, *F*_3,27_ = 7.82, slope *p* = 0.003; see Fig. 2) and increased with wind speed (*m*_*wind*_ = 0.24, slope *p* = 0.004). Full model results are presented in Table 1. Similar relationships existed when the data were subset into vultures that were climbing (slope = -4.63, *F*_1,29_ = 7.57, *p* = 0.01, adj. r^2^ = 0.18) and flying approximately level (slope = -4.52, *F*_1,29_ = 7.51, *p* = 0.01, adj. r^2^ = 0.18), however the relationship between *V*_*med*_ and *ρ* was not significant when the birds were descending (slope = -2.72, *F*_1,29_ = 2.81, *p* = 0.10, adj. r^2^ = 0.06).

**Table 1.**
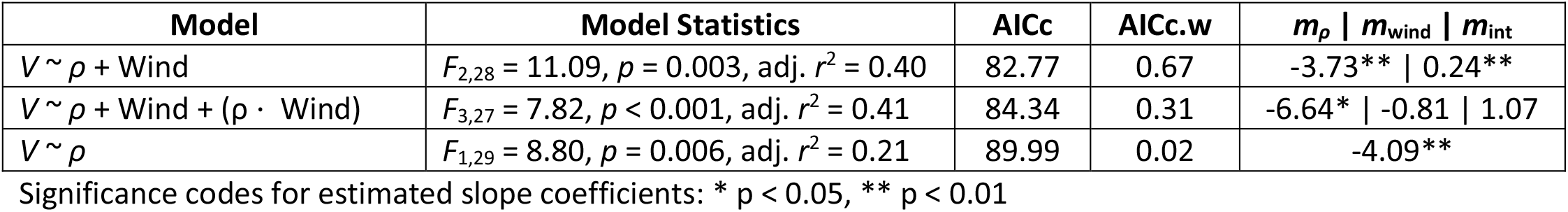
Model selection results

The estimated slope of the relationship between *V*_*med*_ and *ρ* was not statistically distinguishable from the predicted slope (*m*_*pred*_) in the best performing model, which included both *ρ* and wind speed (Wald *χ*^2^ test, *F*_2,28_ = 2.04, p = 0.16), the runner-up model with the wind speed-by-*ρ* interaction (*F*_3,27_ = 0.15, p = 0.70), or the model which included only *ρ* (*F*_1,30_ = 0.67, p = 0.42). Similarly, the slopes of the tests of climbing and level flight were indistinguishable from *m*_*pred*_ (Wald *χ*^2^ test, all p > 0.65).

### Flapping analysis

We were only able to detect flapping in tracks that were in close proximity to the cameras, so we restricted analysis of flapping behavior to tracks that were within 350 m of the roost, and at least 50 m away from it to avoid analyzing landing tracks. Fortunately, 84% of the tracks in the overall dataset fit these criteria (see Fig. 3). There was no relationship between the proportion of flapping in the tracks and *ρ* (OLS regression, *F*_1,29_ = 1.91, *p* = 0.18), however, the proportion of flapping in tracks decreased steeply away from the roost (Fig. 3). There was also no relationship between wind speed and the proportion of tracks where flapping was detected (*p* = 0.59), but the probability of detecting flapping did increase (*p* < 0.01) with wind speed (see Fig. 4).

**Figure 3.**
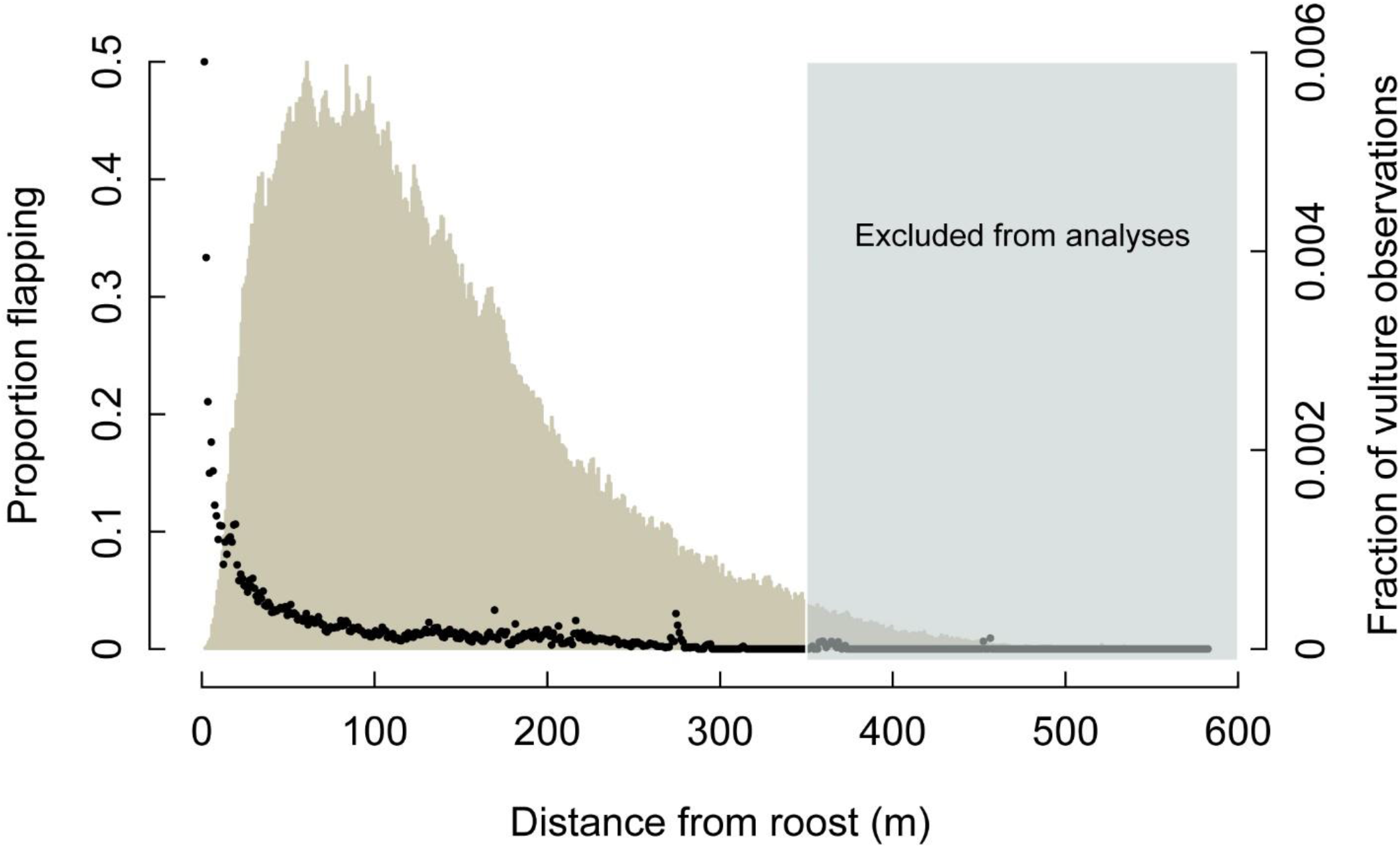
Proportion of detected flapping events, indicated by black points, decreased with distance from the roost trees. The histogram depicts distribution of tracked birds, relative to the roost position. The ability to detect flapping diminished with distance from the cameras, so tracks greater than 350 m from the roost were excluded from further analyses.

**Figure 4.**
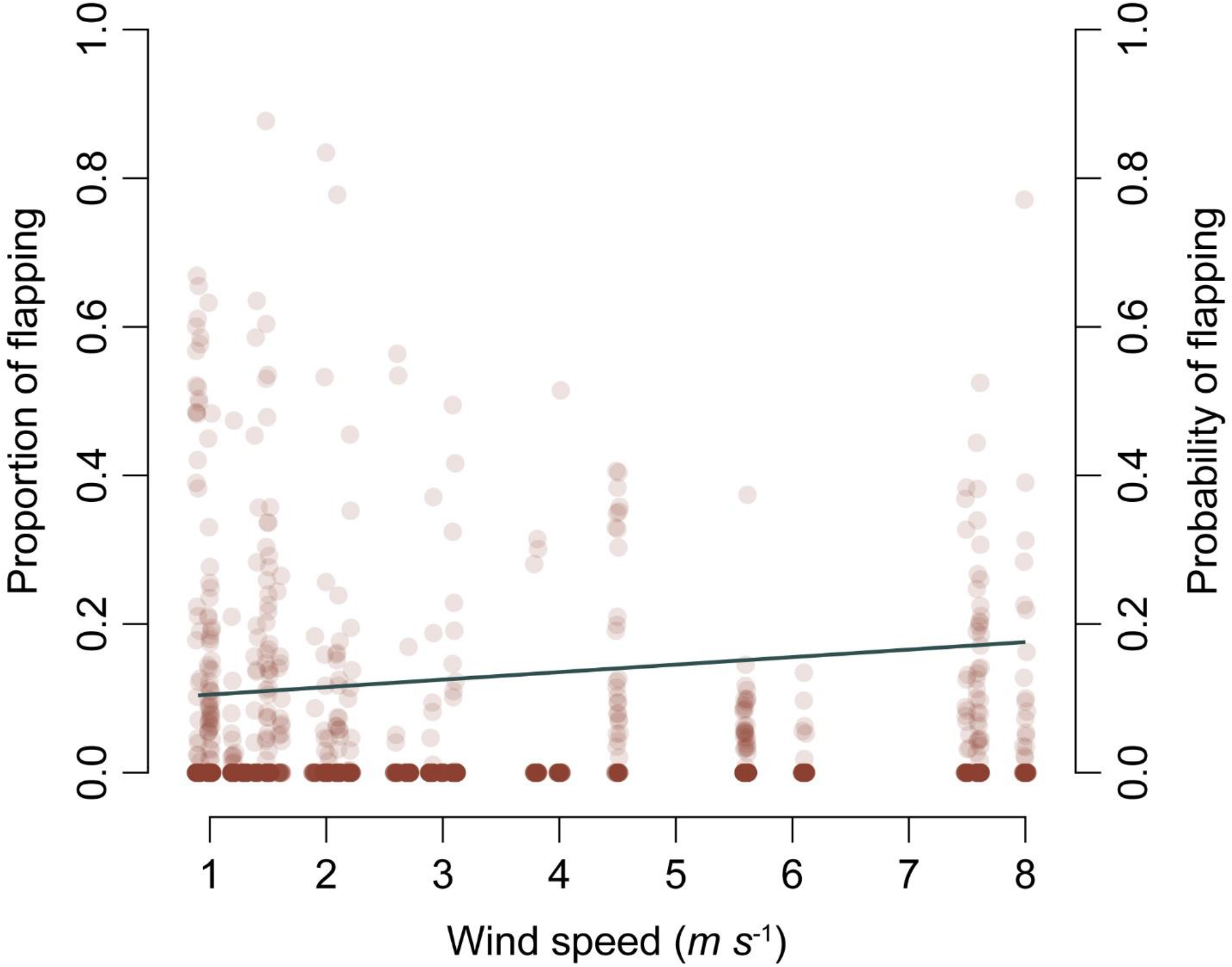
Flapping behavior as a function of wind speed. The proportion of time that birds spent flapping was low across all wind conditions, but increased slightly (though not significantly) with wind speed. Points show individual track values to depict the dispersion of the data, but analyses were conducted on median values for each recording session. The line shows the mean probability of observing flapping in a track, which increased with wind speed.

### Glide polar

Our kinematic analysis of vulture flight tracks produced a glide polar with a minimum sinking speed of 0.74 ms^−1^ at a flight speed of 7.9 ms^−1^ (Fig. 5). Coefficients *k*_1_ and *k*_2_ were both highly significant (*p* < 0.001) and had respective values of -0.0134 and -0.0260 with standard errors of 4.58e^−4^ and 1.00e^−3^. The maximum range speed was 10.3 ms^−1^ with a sinking speed of 0.84 ms^−1^. The maximum lift to drag ratio was 12.2. The mean flight speed in the dataset constructed for fitting the glide polar (i.e. straight, descending glides) was 10.1 ms^−1^, nearly identical to the maximum range speed expressed by the polar. However, the comparison dataset of “gliding turns” had a mean airspeed of 8.0 ms^−1^, nearly identical to the minimum power speed in the glide polar.

**Figure 5.**
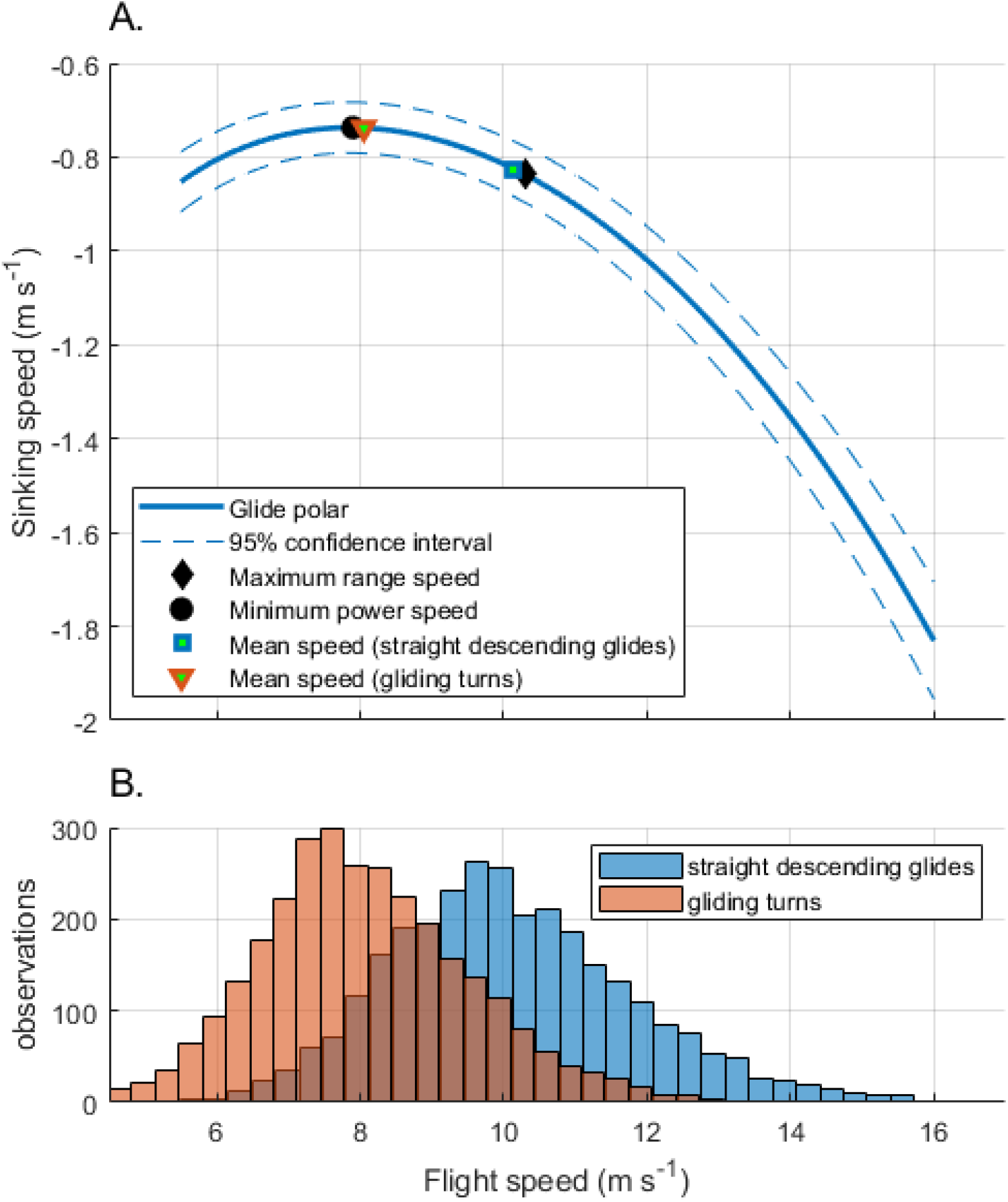
The Turkey vulture glide polar measured from the recorded trajectories. A) shows the glide polar itself, along with a 95% confidence interval, two biomechanically interesting points (black symbols) and the mean flight speed of the vultures during two different types of flight behavior (colored symbols). B) expands the mean speed results for straight glides and gliding turns into histograms to fully illustrate the difference between the two categories. The two distributions have significantly different means (p<0.0001, 2-sample t-test) and these means correspond almost precisely to the two biomechanical optimal for transport over distance (i.e. maximum range speed) during straight glides and minimum descent speed for turning glides. Flying at minimum descent speed will both facilitate the vulture taking advantage of a column of rising air and minimize aerodynamic costs of turning.

## Discussion

### Summary of results

We predicted that median airspeed in vultures flying across a range of elevations and ambient conditions would increase in response to decreasing air density (*ρ*). Based on a simple linear approximation of the relationship between airspeed and *ρ*, we predicted a slope of -5.22 across the sampled range of *ρ*. We found that median vulture airspeed (*V*_*med*_) largely conformed with this prediction, despite a large amount of variation among individual tracks. Furthermore, this relationship was largely invariant with flight behavior; climbing and level tracks also followed the predicted relationship. These results agree with prior observations of Himalayan vultures (*Gyps himalayensis*) tracked via GPS [27].

### Increased airspeed compensates for decreased air density

Drag forces also decrease with air density [32], and is a likely mechanism for the observed increase in airspeed. However, this assumes that birds among the different populations are geometrically similar, having roughly the same wing loading and wing shape, and that morphological disparity (the variation in body shape and size) is roughly equivalent across populations. Local adaptation of wing morphology to high elevation in the sample would likely have manifested in less change in *V*_*med*_ relative to *ρ*, or a difference in the ratio of sinking speed (*V*_*Z*_) relative to horizontal speed (*V*_*XY*_). There was no difference in the relative contribution of horizontal vs. sinking speed in the sample (*p* = 0.99). This, plus the agreement between the predicted and estimated slopes of the *V*_*med*_ vs. *ρ* relationship suggest no localized adaptation in wing morphology or loading. Further, if the increase in airspeed is simply a passive effect related to the reduction of drag, it implies that birds do not alter their flapping behavior [10] or increase their glide angle in low density air. Vultures in the recordings examined here flapped more as they neared their roost trees, perhaps as part of their approach and landing maneuvers. Flapping also increased in response to greater wind speed, but did not vary with *ρ*. Taken together, these results do not indicate any localized behavioral or morphological adaptations to maintain similar airspeed among populations of Turkey vultures residing at different elevations.

Our sample sites, which encompass a 27% reduction in air density, captured the variation of flight conditions that vultures experience during typical flight bouts across their geographic range, but may not reflect extreme conditions that vultures sometimes experience. Vultures have been recorded at altitudes exceeding 1000 m [16], and anecdotal reports suggest that their maximum flight altitudes may be much higher (up to perhaps 6000 m). Air density decreases exponentially as altitude increases, necessitating disproportionally greater airspeed increases to maintain lift as birds climb higher, so it is possible that the relationship that we have described between *V*_*med*_ and *ρ* might change for vultures flying at especially high altitudes, requiring that they modify their flapping behavior or glide angle. Despite some observations of extremely high vulture flight, tracking data suggest that typical vulture flight altitudes are much lower, around 150 m above ground level [16,17], well within our sampled elevation range.

### Wind effects

Wind conditions on our sample days varied from calm to heavy, gusty winds. The best performing model included both air density and wind speed as predictors for vulture airspeed. Our estimate of wind speed is based upon its influence on the measured groundspeeds of the birds flying at different angles relative to the wind (see Methods), and we would expect no lingering relationship between wind speed and airspeed if this correction is consistent across all wind velocities. However, we found a positive relationship between wind speed and airspeed. This relationship was largely driven by the windiest day in our sample, with an average wind speed of 7.8 ms^−1^. The linear regression between median vulture airspeed and estimated wind speed was significant (*p* < 0.01) with that day included but was non-significant when we removed it (*p* = 0.08). Our overall multiple regression model results predicting vulture airspeed were robust to inclusion, or not, of that particularly windy day. We interpret this result as the vultures modifying their flight behavior to maintain forward progress toward their destination in the face of strong headwinds. The wind speed on that day was nearly equal to the minimum power speed (Fig. 5) and thus sufficient to noticeably influence bird airspeed [33]. Despite the wind effect, the relationship between vulture airspeed and air density is statistically indistinguishable from the slope that we predicted based on the elevation and ambient temperature and humidity at our sample sites. Therefore, though the vultures clearly reacted to windy conditions, air density was still the dominant influence on their flight speed.

### Glide polar and gliding performance

Our analysis of the vulture flight trajectories produced a glide polar (Fig. 5) with characteristic minimum power and maximum range speeds of 7.9 and 10.3 ms^−1^, respectively. While no published glide polars exist for Turkey vultures, several exist for Black vultures (*Coragyps atratus*), which are of similar body mass and sympatric with Turkey vultures for much of their range. However, Black vultures have significantly higher wing loading (i.e. smaller wings relative to body mass) compared to Turkey vultures [31,34], and these morphological differences are expected to manifest as differences in flight performance. Specifically, gliding Turkey vultures are expected to have a smaller magnitude minimum sinking speed than Black vultures and achieve that sinking speed at a slower airspeed than Black vultures [35,36]. Our results support these predictions; Parrot [37] studied the gliding flight of a Black vulture in a wind tunnel and reported a minimum sinking speed of approximately 1.1 ms^−1^ at an airspeed of 11.5 ms^−1^, compared with the respective values of 0.74 and 7.9 ms^−1^ for Turkey vultures in this study.

The Turkey vultures studied here were also strikingly proficient at modulating their flight speed toward different biomechanical optima for different behaviors. Straight, descending gliding flight (instantaneous turn radius > 20 m) had a mean equivalent (i.e. air density adjusted) airspeed of 10.1 ms^−1^, similar to the maximum range speed of 10.3 ms^−1^ on the glide polar. Thus, during straight glides the vultures were using a flight speed that maximizes distance traveled per unit energy expended. We did not observe any sustained circling behavior in the vulture flights recorded here, but did record some turning flight, and even instances where the vultures achieved an instantaneous gain in their summed kinetic and potential energy, a result consistent with brief use of rising air currents for energy gain. Birds gaining energy from rising air should fly at an airspeed close to that which minimizes sinking speed thus maximizing the elevation gain from the rising air. Birds turning for other reasons may also slow their airspeed to minimize centripetal acceleration during turning, and flying at the minimum sinking speed minimizes energy losses to induced, parasite, and profile drag leaving more lift available for turning. Both these effects suggest birds engaged in gliding turns should fly near their minimum power speed. In this case, turning Turkey vultures (instantaneous turn radius < 15 m) flew at an airspeed of 8.0 ms^−1^, nearly identical to the minimum power speed of 7.9 ms^−1^ from their glide polar. Thus, Turkey vultures conform surprisingly closely to the separate biomechanical optima for gliding for distance and gliding while turning.

### Concluding remarks

Animals interact with their physical environment to move, forage, migrate, and a host of other functions, and their ability to do so effectively can be limited by physical constraints imposed by their environment. We showed that Turkey vultures respond to a fundamental environmental gradient that could impact their flight performance, the decrease of air density at high elevation by increasing their flight speed and adjust their flight speed to optimize their gliding flight. The tool that we used was developed for field studies of biomechanics [13,14,25,38], but this study also demonstrates that tools from the biomechanics toolchest can be successfully applied to ecological questions.

## Data availability

Vulture tracks and associated metadata are available along with the R and MATLAB data processing scripts used to perform the analyses and generate the figures in this manuscript will be made publicly and permanently available upon manuscript acceptance.

## Acknowledgements

We would like to thank the Orange County Landfill in Chapel Hill, NC, the University of Wyoming in Laramie, WY, and the Alcova Lakeside Marina in Alcova, WY for access to vulture roost sites. Further, we thank Jennifer Heyward, Shanim Patel, and Pranav Khandelwal, and Brenna Hansen for video assistance in Chapel Hill, NC, and Robert Carroll and Braden Godwin for assistance with video collection and site logistics at the Laramie, WY location. William and Judith Rader assisted with video collection at the Alcova, WY site. Dennis Evangelista helped with fieldtrip planning and general advice for data management and processing of vulture tracks.

## Funding

This work was supported by NSF IOS 1253276 to TLH, Office of Naval Research grant N0001410109452 to TLH and 8 others, and by NSF DEB 1737752 to Daniel R. Matute.

## Notes

### Competing Interest Statement

The authors have declared no competing interest.

